# Repeated cocaine reorganizes striatal ensembles along the ventral–dorsal axis

**DOI:** 10.64898/2026.09.15.751852

**Authors:** Michael Z. Leonard, Hannah B. Reiley, Ainoa Konomi Pilkati, Kathryn J. Bjornson, Maxime Chevée, Erin S. Calipari

## Abstract

Drugs of abuse produce lasting adaptations in the striatum. A prominent theory holds that the early reinforcing effects of cocaine preferentially engage the ventral striatum, and repeated cocaine use recruits dorsal striatal systems associated with habitual drug-related behavior. However, this transition has largely been conceptualized on the population level, obscuring how it is implemented within individual neurons and how repeated drug exposure reorganizes drug responsivity within these circuits. Here, we used single-cell calcium imaging to examine ventral and dorsal striatal responses to acute and repeated cocaine exposure in mice. Cocaine broadly suppressed neuronal activity throughout the ventral and dorsal striatum. However, repeated exposure produced opposing changes in cocaine-modulated neurons across the ventral–dorsal axis. The cocaine-activated population was reduced ventrally while expanding dorsally. Unique to dorsal striatum, repeated cocaine engendered two distinct features of activity: 1) Cocaine-activated cells had weak baseline coupling with the surrounding neuronal population and 2) The emergence of a subpopulation of neurons exhibiting regular, slow rhythmic activity in response to cocaine. Thus, repeated cocaine effects reflect a change in how the circuit is organized rather than simply how strongly it is engaged. This organization may allow cocaine to engage patterns of dorsal striatal activity that are largely absent under baseline conditions, creating a circuit state that becomes increasingly specific to the presence of the drug.

## Introduction

The organization of striatal activity changes with experience. During the acquisition of learned behaviors, early engagement of ventral striatal circuits gives way to increasing recruitment of dorsal striatal circuitry as behavior becomes automatic^1^. This normally adaptive reorganization is thought to be co-opted during addiction, such that repeated drug experience increasingly engages dorsal striatal systems that support automatic and habitual behaviors^2–4^. Consistent with this model, chronic cocaine self-administration produces functional changes within the brain that extend from ventral into dorsal striatal regions over experience^5^.

Much of the evidence for progressive dorsal striatal recruitment comes from animals that repeatedly perform actions to obtain cocaine, making it difficult to determine whether dorsal striatal adaptations are a consequence of cocaine exposure itself or of the behavioral experience associated with obtaining the drug. Importantly, cocaine exposure alone can influence this process: noncontingent cocaine facilitates subsequent habit formation and biases associative encoding toward dorsolateral striatum^6,7^ and repeated cocaine injections result in cocaine-evoked ERK activation in dorsal striatum despite the attenuation of most cocaine-responsive signaling pathways in other striatal subregions^8^. These findings raise the possibility that cocaine itself may be sufficient to initiate the reorganization of striatal activity, independent of behavioral control.

However, regional changes in striatal function do not reveal how the underlying drug-responsive neuronal populations are reorganized. Cocaine-responsive neurons – often termed “ensembles”-comprise only a subset of the striatal population, and individual neurons can exhibit heterogeneous responses to the drug. Thus, an apparent change in the engagement of a striatal region as a whole could arise through several distinct forms of population-level reorganization, which could also differ across regions. Here, we directly tested this possibility by tracking single-cell activity across the ventral–dorsal striatal axis during acute and repeated noncontingent cocaine exposure.

## Results and Discussion

We recorded single-cell calcium activity from either ventral or dorsal striatum in freely moving mice during experimenter-administered cocaine (**Fig. 1A**). Responses to cocaine (10 mg/kg) were measured during the initial exposure and again following 10 days of repeated daily administration. Saline administration, alone, had little effect on neuronal activity at either time point (**Fig. S1**).

**Figure 1.**
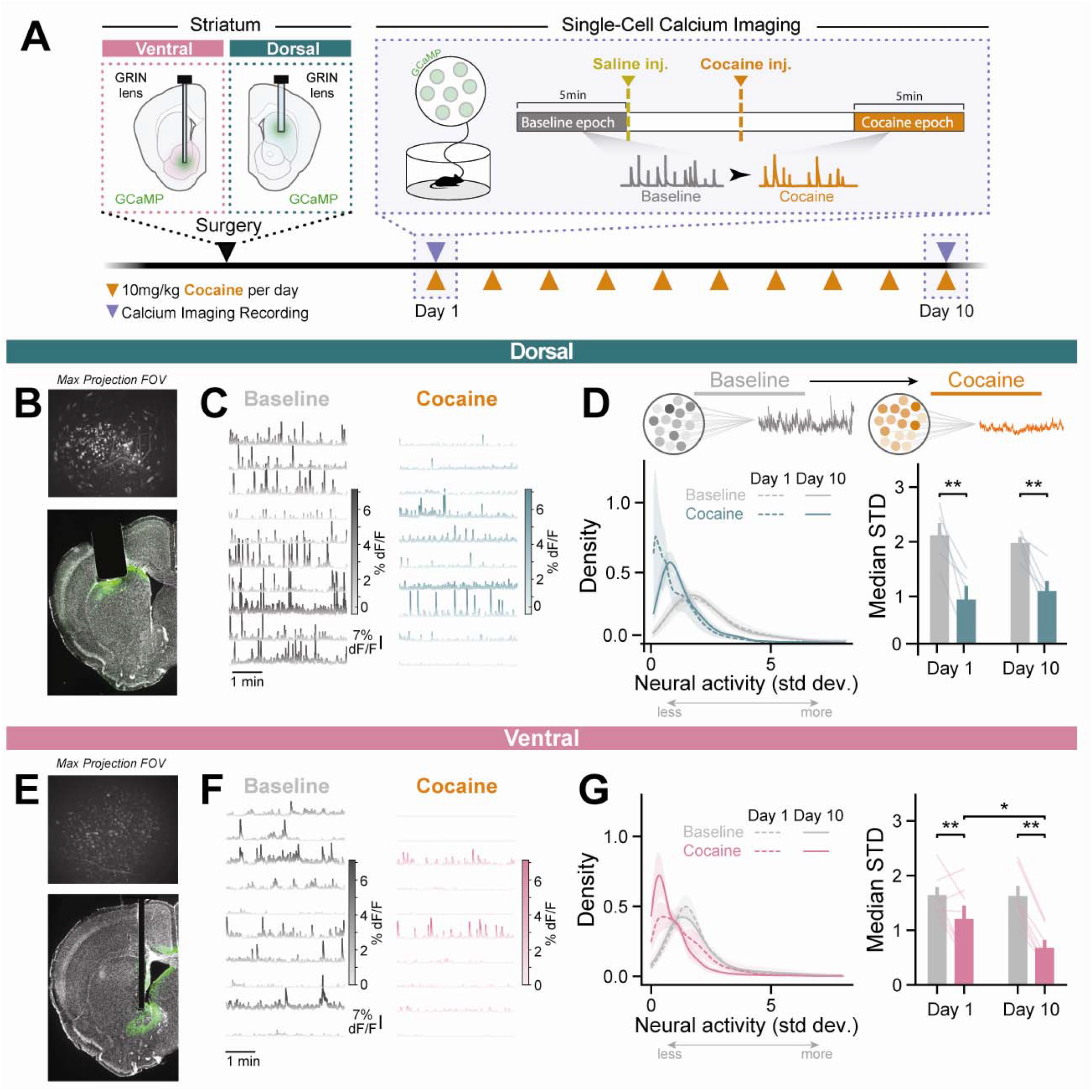
Cocaine broadly suppresses neuronal activity across ventral and dorsal striatum. (**A**) Experimental timeline. (**B**,**E**) Representative histology and fields of view from dorsal (B) and ventral (E) striatum. (**C**,**F**) Representative calcium traces at baseline and following cocaine. (**D**,**G**) Diagram illustrating analysis strategy for population activity (top), distribution *(left)* and median *(right)* of dF/F standard deviation showing cocaine-induced changes in neuronal activity on Days 1 and 10. In dorsal striatum, cocaine reduced activity STD on both days (RM ANOVA: main effect of condition p = 0.015; main effect of day p = 0.86; interaction p = 0.36; N = 5 mice). In ventral striatum, cocaine similarly reduced activity STD, with a differential effect across days (RM ANOVA: main effect of condition p = 0.0029; main effect of day p = 0.034; interaction p = 0.042; N = 7 mice).

Cocaine reduced activity across both dorsal (**Fig. 1C,D**) and ventral striatum (**Fig. 1F,G**), despite considerable heterogeneity in the magnitude and direction of individual-cell responses. This population-level suppression remained prominent following repeated cocaine exposure (**Fig. 1D,G**). In ventral striatum, cocaine-induced suppression was enhanced after repeated exposure, whereas the magnitude of suppression remained comparatively stable dorsally. This widespread suppression is consistent with electrophysiological and metabolic studies identifying inhibition as a principal accumbal response to cocaine^5,9^, and showing that repeated cocaine experience can potentiate population hypoactivity^10,11^. We therefore asked whether repeated cocaine differentially reorganized the minority of neurons that *increased* their activity in response to cocaine.

Despite this widespread suppression, a small population of neurons increased activity following cocaine, and repeated exposure had opposing effects on these cocaine-responsive populations across striatal regions (**Fig. 2**; region × day interaction, *p* = 0.024). In ventral striatum, the proportion of cocaine-activated neurons decreased with repeated exposure, whereas the cocaine-activated population expanded in dorsal striatum. Repeated cocaine can differentially restructure accumbal subpopulations, increasing recruitment of cocaine-activated D1-MSNs while enhancing suppression among D2-MSNs during sensitization^12^. Although our recordings do not distinguish MSN subtype, the net contraction of the ventral cocaine-activated population is consistent with evidence that repeated cocaine progressively refines cocaine-responsive ensembles in the ventral striatum, where increasingly selective neuronal populations can acquire greater influence over cocaine-associated behaviors^11,13^. Thus, reduced ensemble size may not reflect diminished ventral striatal involvement, but may instead reflect increasing selectivity within the drug-responsive population. Together, these opposing adaptations reveal that repeated cocaine does not uniformly expand or contract drug-responsive ensembles, but reorganizes cocaine responsivity across the ventral–dorsal striatal axis: refining the responsive population ventrally while progressively recruiting dorsal striatal circuitry.

**Figure 2.**
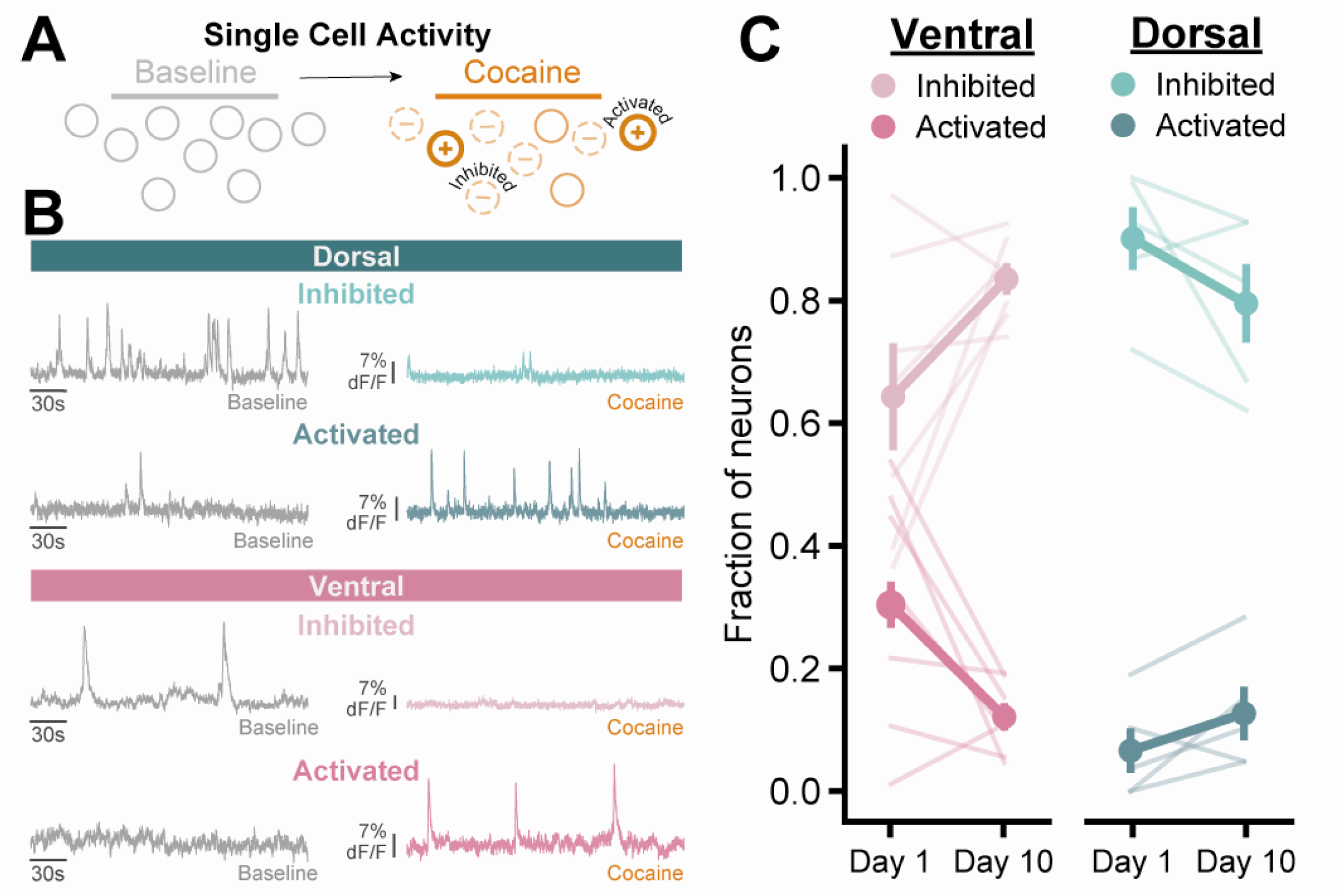
Repeated cocaine differentially reorganizes cocaine-responsive ensembles across ventral and dorsal striatum. (**A**) Diagram illustrating analysis strategy for cocaine-response phenotypes and (**B**) representative cocaine-inhibited and -activated neurons in ventral and dorsal striatum. (**C**) Proportion of cocaine-inhibited and activated neurons on Days 1 and 10. Repeated cocaine differentially reorganized ensemble recruitment across regions, with distinct proportions of inhibited (mixed ANOVA interaction: p = 0.026) and activated (mixed ANOVA interaction: p = 0.024) neurons. Descriptively, the activated population decreased from 30.6 ± 7.5% to 12.3 ± 2.2% in ventral striatum, while increasing from 6.6 ± 3.6% to 12.6 ± 4.4% in dorsal striatum. The inhibited population showed the opposite pattern, increasing from 64.6 ± 8.7% to 83.7 ± 2.5% ventrally and decreasing from 90.1 ± 5.1% to 79.5 ± 6.4% dorsally. Data represented as means ± SEM.

We next asked whether the dorsal neurons recruited by cocaine after repeated exposure were functionally distinct *before the drug was administered*. To test this, we identified neurons based on their response to cocaine and then examined their relationship to the surrounding population during the preceding baseline period, before cocaine had been administered (**Fig. 3A**). By Day 10, cocaine-activated neurons in dorsal striatum were less correlated with other simultaneously recorded neurons at baseline than those identified during the cocaine exposure on Day 1 (**Fig. 3A,B**). Thus, repeated cocaine activated dorsal striatal neurons that were uncoupled from ongoing population activity, rather than preferentially engaging neurons already strongly correlated with other neurons in the surrounding network. This difference was not observed in the ventral striatum.

**Figure 3.**
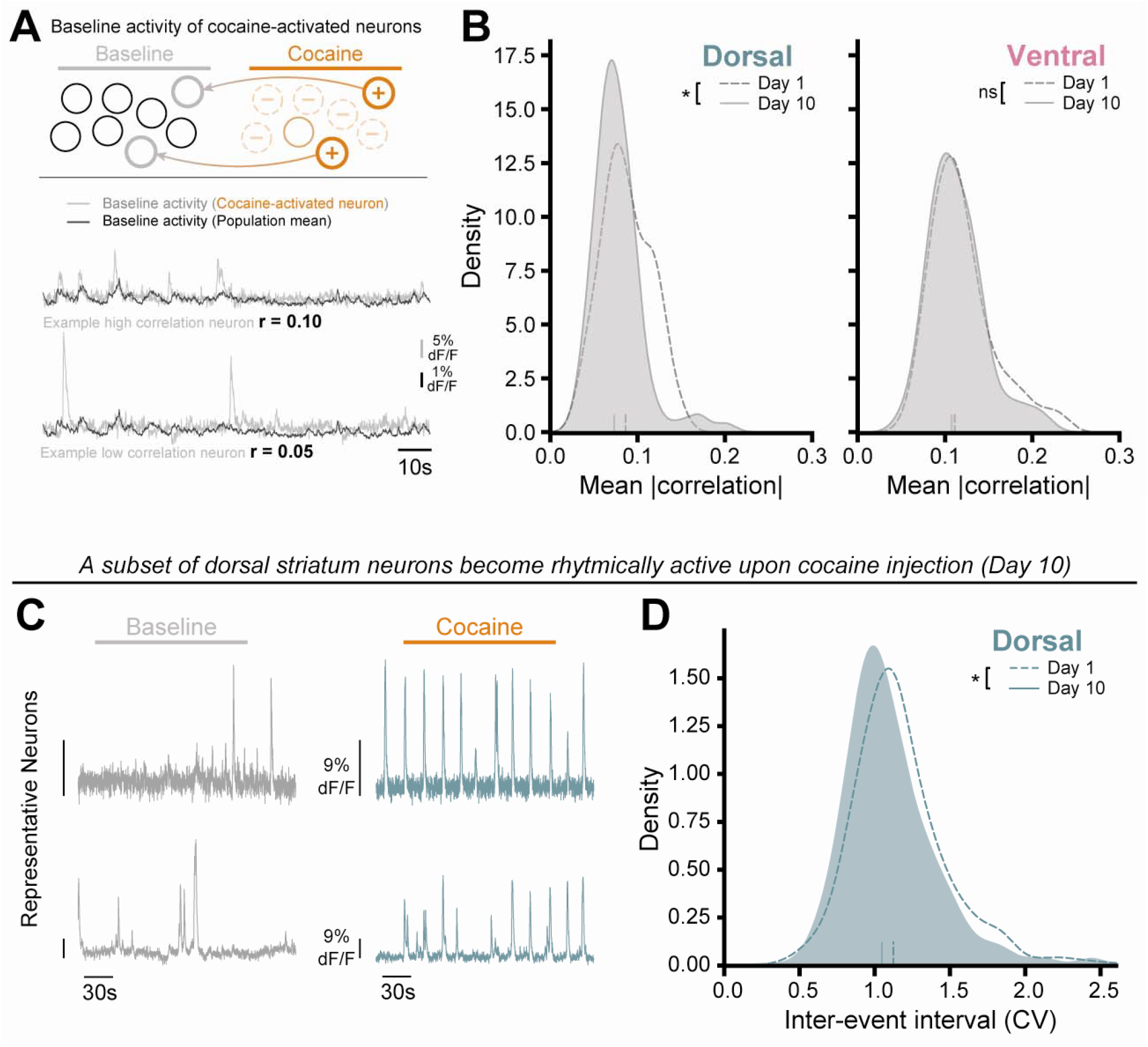
Repeated cocaine recruits weakly population-coupled dorsal neurons and promotes slow rhythmic activity. (**A**) Diagram illustrating analysis strategy and example neuron traces showing different correlations to population mean activity. (**B**) Distribution of the mean pairwise correlations for each cocaine-activated neuron with all other simultaneously recorded neurons on Days 1 and 10 in dorsal and ventral striatum. Repeated cocaine reduced population coupling selectively in dorsal cocaine-activated neurons, at baseline (Day 1: n = 52 cells; Day 10: n = 87 cells; two-sample KS test with permutation, KS = 0.236, p = 0.035). No significant change was observed in ventral striatum (Day 1: n = 111 cells; Day 10: n = 68 cells; KS = 0.126, p = 0.451). (**C**) Representative calcium traces showing increased regularity of cocaine-evoked activity in dorsal striatum following repeated cocaine. (**D**) Distribution of the coefficient of variation of inter-event intervals, showing increased regularity of cocaine-evoked activity following repeated cocaine. In dorsal striatum, cocaine-evoked activity became significantly more regular from Day 1 to Day 10 (Day 1: n = 497 cells; Day 10: n = 505 cells; two-sample KS test with permutation, KS = 0.154, p < 0.001).

The dorsal striatal population recruited after repeated cocaine was also distinguished by its activity in the presence of the drug. By Day 10, a subset of dorsal neurons exhibited slow, rhythmic activity following cocaine administration, with calcium events occurring at more regular intervals than during the initial cocaine exposure (**Fig. 3D–F**). Slow oscillations on similar timescales have been described throughout basal ganglia circuitry and are strongly regulated by dopamine, which can increase their regularity^14^. Such slow basal ganglia dynamics have been proposed to organize extended motor sequences and, when exaggerated by dopaminergic stimulation, may promote persistent patterns of behavior that are less responsive to ongoing environmental feedback. Thus, repeated cocaine recruited dorsal striatal neurons distinguished not only by their relationship to ongoing population activity before drug administration, but also by the emergence of highly regular activity in the presence of cocaine.

These findings have implications for how progressive dorsal striatal recruitment is interpreted in addiction. The ventral-to-dorsal transition is often conceptualized as increasing engagement of dorsal striatal circuitry as drug-seeking becomes habitual. Our findings demonstrate that repeated cocaine does not increase dorsal striatal activity globally, but instead changes how neurons respond to drug over time. Moreover, this redistribution occurred without an instrumental action or drug-seeking experience, indicating that dorsal striatal recruitment need not arise as a consequence of explicit drug-reinforcement. Instead, repeated pharmacological exposure may begin reorganizing striatal representations before these behaviors develop, potentially biasing subsequent learning toward dorsal striatal control. The preferential recruitment of relatively weakly coupled dorsal striatal neurons also raises the possibility that repeated cocaine alters the rules governing ensemble allocation, progressively incorporating neurons outside dominant ongoing population dynamics, with unique response properties, into the drug-modulated population.

## Materials and Methods

Calcium activity (AAV1.CaMk2a.GCaMP6m) was recorded in the dorsal or ventral striatum of male and female mice using an Inscopix nVista miniature fluorescence microscope in response to acute and repeated cocaine (10 mg/kg/day). Detailed description of the methods is provided in *Extended Methods*.

## Acknowledgments

We thank members of the Calipari lab for thoughtful discussions of the manuscript. ESC was supported by National Institutes of Health (NIH) grants R01DA052317, R01AA030931 and P60AA031124; MC by NIH grants K99DA060930, and a BBRF NARSAD Young Investigator Grant #32921; MZL by NIH grant F32DA060662; HBR by NIH grant F32DA061606; KJB by NIH grant F32AA032415; AKP by the Independent Research Fund Denmark, DFF.

## Author contributions

MZL, MC, HBR, and ESC conceived and designed the experiments; MZL, MC, HBR, and AKP performed miniscope recordings; MC and MZL performed histology; MC wrote code and analyzed the data, MZL, HBR, MC, KJB and ESC wrote the manuscript with input and approval from all authors.

## Competing Interest Statement

The authors declare no competing interests.

---

**Figure S1.**
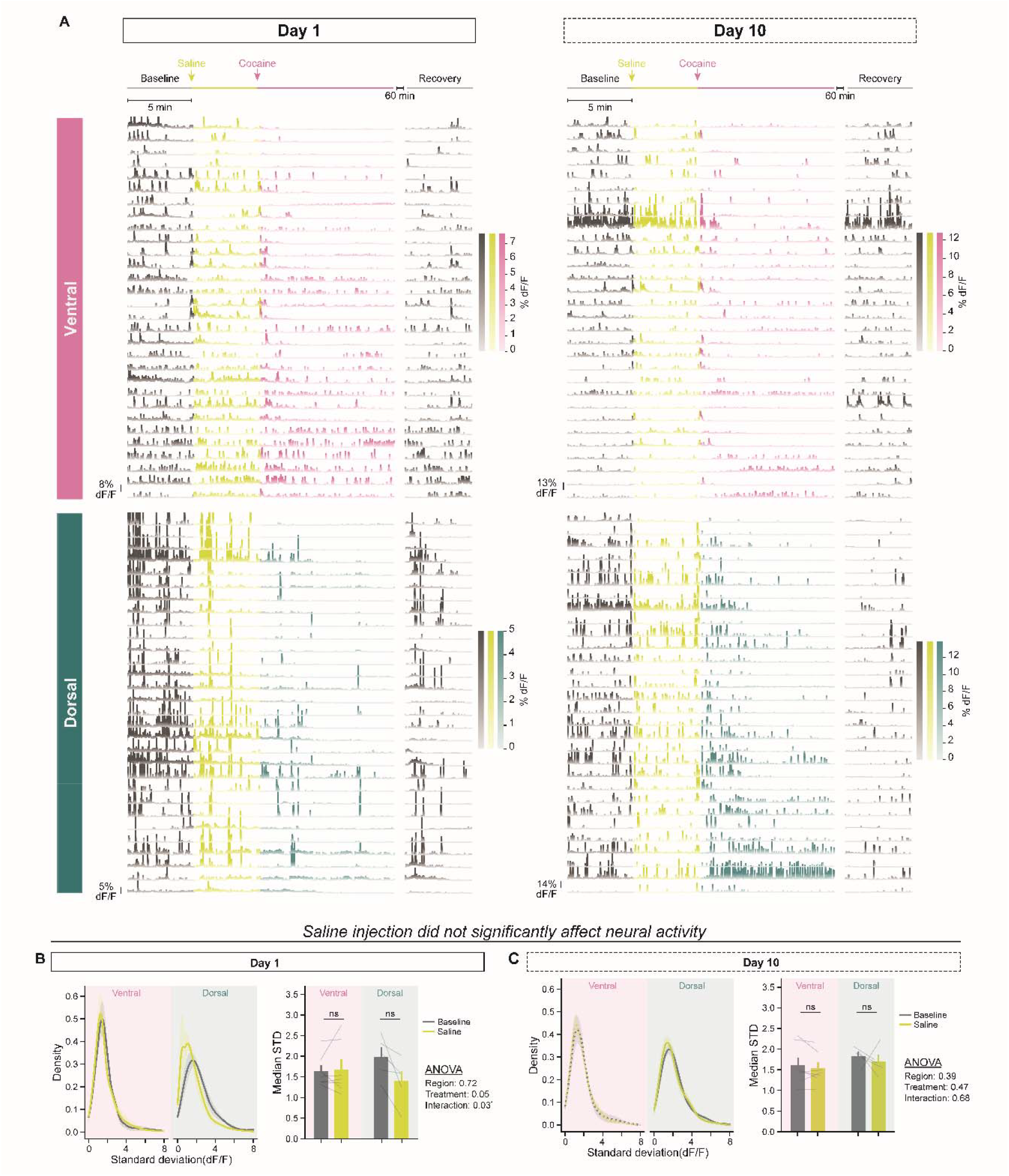
Saline administration does not significantly modulate striatal neuronal activity. (**A**) Representative calcium traces from ventral and dorsal striatum during baseline, saline, cocaine, and recovery periods on Days 1 and 10. **(B, C)** Distribution (left) and median (right) of dF/F standard deviation on Day 1 (B) and Day 10 (C) in ventral and dorsal striatum following saline administration. On Day 1, saline did not significantly change activity STD in either dorsal (Baseline: 1.989 ± 0.227; Saline: 1.401 ± 0.245; Wilcoxon signed-rank, p = 0.063) or ventral striatum (Baseline: 1.640 ± 0.145; Saline: 1.682 ± 0.238; Wilcoxon signed-rank, p = 0.938). On Day 10, saline similarly had no significant effect in either dorsal (Baseline: 1.847 ± 0.104; Saline: 1.727 ± 0.151; Wilcoxon signed-rank, p = 0.625) or ventral striatum (Baseline: 1.625 ± 0.185; Saline: 1.553 ± 0.147; Wilcoxon signed-rank, p = 0.375). n = 5 dorsal, 7 ventral mice.

## Extended Methods

### Subjects

Experiments were approved by the Institutional Animal Care and Use Committee of Vanderbilt University and conducted according to the National Institutes of Health guidelines for animal care and use. Twelve adult (16 to 36-week-old) mice were used for this study. C57BL/6J mice (6 males and 6 females) were acquired from Jackson Laboratory (Bar Harbor, ME; SN: 000,664) and maintained on an 8am/8pm 12-h reverse light cycle. Mice had unlimited access to food and water. Experiments were performed during the dark phase.

### Viral injection and GRIN lens implantation

To monitor neuronal activity in the ventral and dorsal striatum, we injected a calcium indicator (AAV1.CaMk2a.GCaMP6m.WPRE.SV40, Inscopix) into the ventral striatum (coordinates relative to Bregma: [AP: +1.40, ML: ± 0.80, DV: −4.30], N=7 mice) and dorsal striatum (coordinates relative to Bregma: [AP: 1.00, ML: ±1.40, DV: −2.50], N=5 mice) in separate cohorts of mice. Thirty minutes prior to surgery, mice were injected 1mg/kg ketoprofen subcutaneously. Mice were anesthetized with isoflurane (induction 2.5%, maintenance 1–2%) and placed on a stereotaxic frame (Kopf Instruments, Los Angeles, CA). Ophthalmic ointment was applied to the eyes and the scalp was shaved and cleaned using aseptic technique. A small craniotomy was made above the target coordinates using a dental drill, and a glass pipette (10–20 µm tip diameter) loaded with virus was lowered to the injection site. 300 nL of virus were pressure-injected and the pipette was left in place for 5–10 minutes before being slowly withdrawn.

For the dorsal striatum, following viral injection, a column of tissue above the injection site was aspirated to a depth of approximately −2.0 mm using a blunt needle attached to vacuum suction, with constant irrigation with sterile saline. A gradient-refractive index (GRIN) lens integrated within a miniscope baseplate (1.0 mm diameter, 4.0 mm length, Inscopix) was then slowly lowered into the aspirated cavity to a final depth of −2.40 mm relative to Bregma and secured to the skull using a layer of C&B Metabond (SKU: S380, Parkell, NY) followed by a layer of dental cement. A similar procedure was used for ventral striatal implants, whereby a blunt needle was lowered to a depth of −4.0mm to provide a tract for the GRIN lens (0.66mm diameter, 7.3mm length, Inscopix), but no tissue was aspirated. A protective cap made of quick-kast silicone elastomer (World Precision Instruments) was placed over the lens to protect it. Mice were allowed to recover for 4 weeks to permit viral expression and resolution of surgery-related inflammation.

Animals were monitored for signs of distress and administered analgesic for 3 days following each surgery.

### Cocaine injections

Cocaine injections were performed as previously described^11^. Briefly, mice received cocaine (10 mg/kg, i.p.) once daily for 10 consecutive days in the same experimental chamber. Calcium imaging was performed during cocaine exposure on Days 1 and 10. On Days 2–9, mice received cocaine without the miniscope attached.

### Miniscope imaging

Calcium activity was recorded using an Inscopix nVista miniature fluorescence microscope, attached to the implanted baseplate immediately prior to each recording session. The mounted blue LED was used to excite GCaMP, and emitted fluorescence was recorded at 20 Hz using the Inscopix Data Acquisition Software. On Days 1 and 10, mice were habituated to the experimental chamber for 10min before imaging. Recordings began with a 5min baseline period, followed by an injection of saline (i.p.). Five minutes later, mice received cocaine (10 mg/kg, i.p.), and recording continued for an additional 10min, for a total recording duration of 20min.

### Data pre-processing

Raw imaging videos were spatially downsampled (2x), spatial bandpass filtered and motion-corrected using the Inscopix Data Processing software. Motion-corrected videos were then processed using constrained non-negative matrix factorization for endoscopic data (CNMF-E) to extract individual putative neurons and their corresponding calcium traces. Each cell was inspected manually and accepted/rejected from the data set. Raw calcium traces were denoised using an OASIS deconvolution algorithm prior to analysis.

### Data analysis

#### Quantification of neuronal activity

For each region, the standard deviation (STD) of denoised activity was compared across day (Day 1, Day 10) and condition (Baseline, Cocaine) using a two-way repeated-measures ANOVA (subject = animal).

#### Defining cocaine-modulated neurons

For each neuron, we calculated a modulation index as Index = (STD cocaine – STD baseline) / (STD cocaine + STD baseline) A neuron was classified as “activated” following cocaine injection if the index was greater than 0.05, and classified as “inhibited” if the modulation index was less than −0.05.

#### Fraction of activated neurons

The fraction of activated cells was calculated per animal per session as the number of cells meeting this criterion divided by the total number of cells recorded in that session.

#### Fraction of activated/inhibited neurons

The fraction of activated and inhibited neurons (as defined above) was calculated per animal per session. Region (Dorsal, Ventral) × day (Day 1, Day 10) effects were assessed using a mixed-design ANOVA (region as a between-subjects factor, day as a within-subjects factor), separately for activated and inhibited fractions.

#### Population coupling

For each cocaine-activated neuron (defined above), we computed the Pearson correlation coefficient between its Baseline-period activity and that of every other simultaneously recorded neuron in the same session, then averaged the absolute value of these pairwise correlations to yield a single population-coupling value per cell. Distributions of population-coupling values were compared between Day 1 and Day 10 within each region using a two-sample Kolmogorov-Smirnov (KS) test; significance was assessed via a permutation test (1,000 iterations, shuffling Day 1/Day 10 labels across pooled cells).

#### Rhythmic activity

Calcium transients were detected on z-scored, denoised traces using a peak-detection algorithm (minimum peak height = 1 SD, minimum peak prominence = 1 SD, minimum inter-peak distance = 0.5 s). For cells with at least 5 detected events, the coefficient of variation (CV) of inter-event intervals (IEI) was calculated as a measure of firing regularity, with lower CV indicating more regular (rhythmic) activity. Distributions of IEI CV were compared between Day 1 and Day 10 for cocaine-evoked activity in each region using the same permutation approach described above.

### Histology

Following recordings, mice were transcardially perfused with 0.01 M phosphate buffered saline (PBS) followed by 4% paraformaldehyde (PFA) solution. Brains were dissected and post-fixed overnight at 4°C in 4% PFA. They were then rinsed 3x in PBS and transferred to a 30% sucrose PBS solution until they sank to the bottom of the tube. Once equilibrated, brains were mounted and frozen onto a sliding microtome (Leica SM2010R) where 60 µm sections were cut. Sections were processed for immunohistochemistry (1:2000 Chicken anti-GFP, GFP-1020, Aves, RRID:AB_10000240 followed by 1:1000 Alexa Fluor 488 AffiniPure Donkey anti-Chicken IgY (IgG) (H+L) antibody, Jackson ImmunoResearch Laboratories Inc, 703-545-155, RRID:AB_2340375). Specifically, sections were incubated in blocking solution (5% Bovine Serum Albumin, 0.3% Triton X-100 in PBS) for 1hr on a rocker, followed by overnight incubation at 4 degrees Celsius with the primary antibodies (diluted in blocking solution). The following day, sections were rinsed 3 times for 10 min in PBS and incubated for 2hrs at room temperature with secondary antibodies diluted in blocking solution. Sections were then rinsed 3 times for 10 min before mounting. Sections were mounted using Prolong Gold antifade with DAPI (Invitrogen) and visualized on an epifluorescence microscope (BZ-X 710, Keyence) using a 4x objective.

### Statistical Analyses

All statistical analyses are described in detail above and were performed using custom code in Python (v3.6.13). The SciPy package (v1.5.3) was used to perform statistical tests and the Pingouin package (v0.3.12) was used to perform two-way and three-way ANOVAs. All data are reported as Mean ± SEM.

## Notes

### Competing Interest Statement

The authors have declared no competing interest.

## References

1. Yin, H. H. et al. Dynamic reorganization of striatal circuits during the acquisition and consolidation of a skill. Nat Neurosci 12, 333–341 (2009).

2. Ito, R., Dalley, J. W., Robbins, T. W. & Everitt, B. J. Dopamine Release in the Dorsal Striatum during Cocaine-Seeking Behavior under the Control of a Drug-Associated Cue. J. Neurosci. 22, 6247–6253 (2002).

3. Belin, D. & Everitt, B. J. Cocaine Seeking Habits Depend upon Dopamine-Dependent Serial Connectivity Linking the Ventral with the Dorsal Striatum. Neuron 57, 432–441 (2008).

4. Everitt, B. J. & Robbins, T. W. Drug Addiction: Updating Actions to Habits to Compulsions Ten Years On. Annu. Rev. Psychol. 67, 23–50 (2016).

5. Porrino, L. J., Lyons, D., Smith, H. R., Daunais, J. B. & Nader, M. A. Cocaine self-administration produces a progressive involvement of limbic, association, and sensorimotor striatal domains. J Neurosci 24, 3554–3562 (2004).

6. Nelson, A. & Killcross, A. S. Amphetamine Exposure Enhances Habit Formation. Journal of Neuroscience 26, 3805–3812 (2006).

7. Takahashi, Y., Roesch, M. R., Stalnaker, T. A. & Schoenbaum, G. Cocaine exposure shifts the balance of associative encoding from ventral to dorsolateral striatum. Front Integr Neurosci 1, ihpa51247 (2007).

8. Bertran-Gonzalez, J. et al. Opposing Patterns of Signaling Activation in Dopamine D_1_ and D_2_ Receptor-Expressing Striatal Neurons in Response to Cocaine and Haloperidol. J. Neurosci. 28, 5671–5685 (2008).

9. Risinger, R. C. et al. Neural correlates of high and craving during cocaine self-administration using BOLD fMRI. NeuroImage 26, 1097–1108 (2005).

10. Peoples, L. L., J. Uzwiak A., Guyette, F. X. & West, M. O. Tonic inhibition of single nucleus accumbens neurons in the rat: a predominant but not exclusive firing pattern induced by cocaine self-administration sessions. Neuroscience 86, 13–22 (1998).

11. Thibeault, K. C. et al. A Cocaine-Activated Ensemble Exerts Increased Control Over Behavior While Decreasing in Size. Biological Psychiatry 97, 590–601 (2025).

12. Van Zessen, R. et al. Dynamic dichotomy of accumbal population activity underlies cocaine sensitization. eLife 10, e66048 (2021).

13. Salery, M. et al. Cocaine-context memories are transcriptionally encoded in nucleus accumbens Arc ensembles. Nat Commun 16, 6084 (2025).

14. Ruskin, D. N. et al. Multisecond Oscillations in Firing Rate in the Basal Ganglia: Robust Modulation by Dopamine Receptor Activation and Anesthesia. Journal of Neurophysiology 81, 2046–2055 (1999).

