## Supplementary material for "Repeated cocaine reorganizes striatal ensembles along the ventral–dorsal axis": Figure S1

### Supplemental Figures

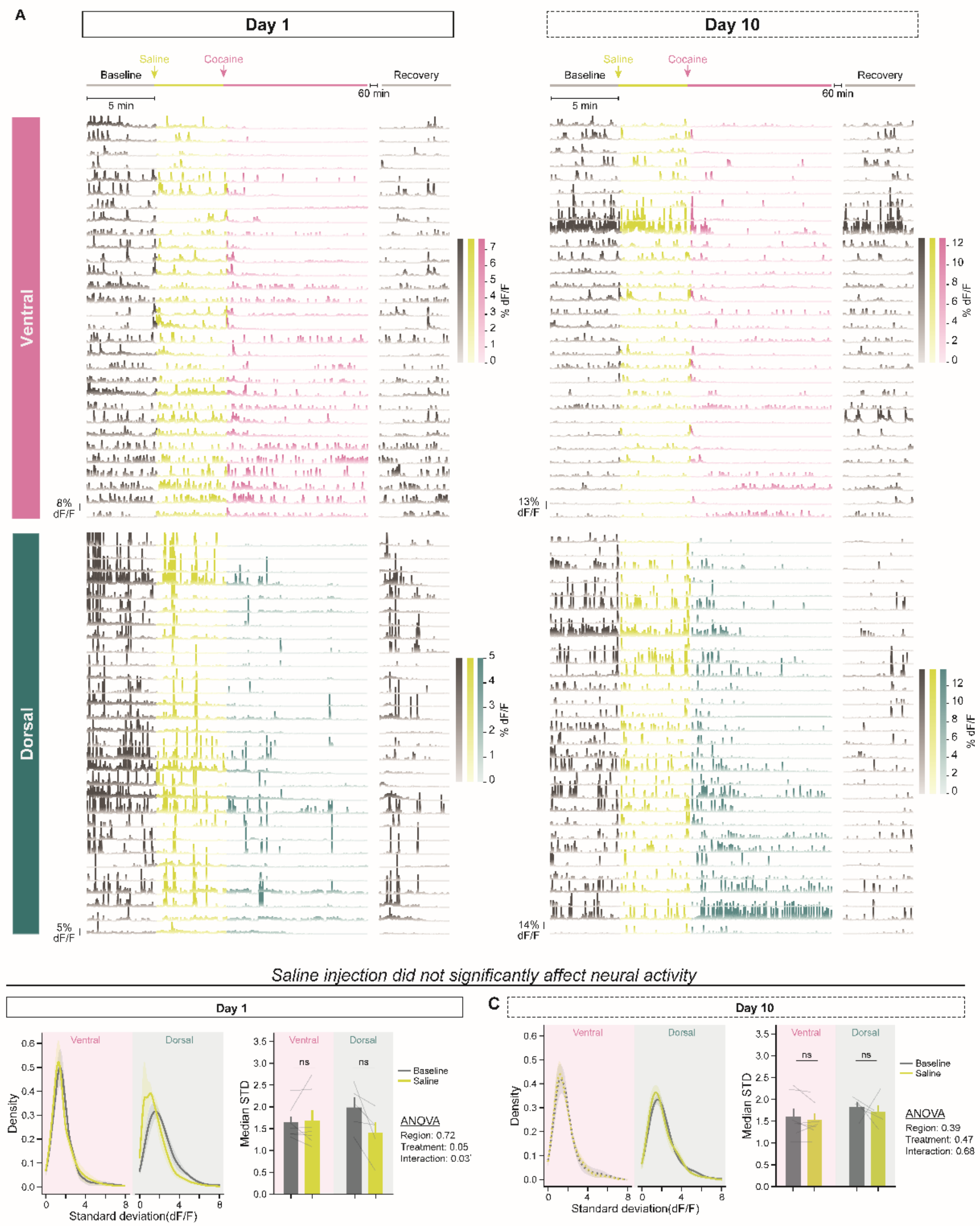

**Figure S1.** *Saline administration does not significantly modulate striatal neuronal activity.*

(**A**) Representative calcium traces from ventral and dorsal striatum during baseline, saline, cocaine, and recovery periods on Days 1 and 10. **(B, C)** Distribution (left) and median (right) of dF/F standard deviation on Day 1 (B) and Day 10 (C) in ventral and dorsal striatum following saline administration. On Day 1, saline did not significantly change activity STD in either dorsal (Baseline: 1.989 ± 0.227; Saline: 1.401 ± 0.245; Wilcoxon signed-rank, $p = 0.063$) or ventral striatum (Baseline: 1.640 ± 0.145; Saline: 1.682 ± 0.238; Wilcoxon signed-rank, $p = 0.938$). On Day 10, saline similarly had no significant effect in either dorsal (Baseline: 1.847 ± 0.104; Saline: 1.727 ± 0.151; Wilcoxon signed-rank, $p = 0.625$) or ventral striatum (Baseline: 1.625 ± 0.185; Saline: 1.553 ± 0.147; Wilcoxon signed-rank, $p = 0.375$). n = 5 dorsal, 7 ventral mice.
